# Increased structural connectivity in high schizotypy

**DOI:** 10.1101/2022.05.12.491533

**Authors:** Eirini Messaritaki, Sonya Foley, Kali Barawi, Ulrich Ettinger, Derek K Jones

**Affiliations:** Cardiff University Brain Research Imaging Centre (CUBRIC), School of Psychology, Cardiff University, Maindy Road, Cardiff, CF24 4HQ, UK; School of Medicine, Cardiff University, Cardiff, UK; Department of Psychology, University of Bonn, Bonn, Germany

## Abstract

The link between brain structural connectivity and schizotypy was explored in two healthy-participant cohorts, collected at two different neuroimaging centres, comprising 140 and 115 participants respectively. The participants completed the Schizotypal Personality Questionnaire (SPQ), through which their schizotypy scores were calculated. Diffusion-MRI data were used to perform tractography and to generate the structural brain networks of the participants. The edges of the networks were weighted with the inverse radial diffusivity. Graph theoretical metrics of the default-mode, sensorimotor, visual and auditory subnetworks were derived and their correlation coefficients with the schizotypy scores were calculated. To the best of our knowledge, this is the first time that graph theoretical measures of structural brain networks are investigated in relation to schizotypy.

A positive correlation was found between the schizotypy score and the mean node degree and mean clustering coefficient of the sensorimotor and the default-mode subnetworks. The nodes driving these correlations were the right postcentral gyrus, the left paracentral lobule, the right superior frontal gyrus, the left parahippocampal gyrus and the bilateral precuneus, i.e., nodes that exhibit compromised functional connectivity in schizophrenia. Implications for schizophrenia and schizotypy are discussed.

## 1. Introduction

Schizophrenia is a serious psychiatric disorder of unknown aetiology, while schizotypy encompasses a set of stable personality traits that are thought to reflect the subclinical expression of schizophrenia. Specifically, people with high levels of schizotypy show cognitive and neural patterns resembling the deficits observed in schizophrenia [1, 2]. Studying schizotypy is, therefore, important for a number of reasons. Firstly, given that partly overlapping factors underlie both conditions [51], studying schizotypy can shed light on the aetiology of schizophrenia. In addition, people with high schizotypy suffer from lower social, educational and professional levels of functioning, and high levels of distress [52, 53]; therefore, studying schizotypy can help us better understand these problems and develop interventions to ameliorate them. Moreover, studying schizotypy can reveal characteristics (in the brains of healthy high-schizotypy participants) which protect or compensate against schizophrenia; those characteristics may be used to inform the development of novel treatments [1, 2]. Finally, the study of schizotypy is free from any confounds that stem from pharmacological interventions, and as such it is of value to both clinical and nonclinical research.

Given this similarity of schizophrenia and schizotypy, in building our aims for this study, we looked to the topological properties of the structural and functional brain networks of schizophrenia patients.

The functional brain networks of schizophrenia patients are compromised and generally exhibit lower connectivity and less efficient organisation, with few exceptions, compared to healthy individuals. For example, a meta-analysis of 13 studies of functional whole-brain networks, in which the functional connectivity was measured via functional MRI (fMRI), magnetoencephalography (MEG) or electroencephalography (EEG), showed significantly lower small-worldness and measures of local organisation [3]. Additional studies that were not included in this meta-analysis corroborate the findings. For example, fMRI-measured functional connectivity was lower within the default-mode network in schizophrenia patients compared to healthy controls [4]. Schizophrenia patients showed lower activation compared to healthy controls in the lateral frontal cortex bilaterally, in the left basal ganglia and the cerebellum [5]. The lateral default-mode network exhibited lower functional connectivity with the sensorimotor cortex, but higher connectivity with association areas in schizophrenia patients [6]. The anterior and posterior default-mode networks exhibited higher and lower connectivity with the right control and lateral visual networks respectively, in schizophrenia patients [6]. Analysis of dynamic functional connectivity showed that schizophrenia patients activated the default-mode network less frequently than healthy controls, but each activation lasted 4-5 s longer than in the healthy controls [7]. Altered functional connectivity was observed in sensorimotor, visual, auditory, social, high level cognitive and motor processing nodes in schizophrenia patients compared to controls [8, 9, 10], with most of the alterations indicating lower functional connectivity. Schizophrenia patients exhibited lower clustering coefficient, global efficiency and local efficiency, and higher characteristic path length of functional networks [11]. Finally, compared to healthy controls, significant hypoconnectivities were observed between seed regions and the areas in the auditory, default mode, self-referential and somatomotor network in schizophrenia patients [12].

The structural brain networks of schizophrenia patients have been shown predominantly to have a less central and less efficient organisation compared to those of healthy controls, but there are some exceptions. For example, the global efficiency, local efficiency, clustering coefficient, and mean connectivity strength of structural networks weighted by the connectivity probability were significantly lower in schizophrenia patients compared to controls, while the mean betweenness centrality was significantly higher [13]. Significantly longer path lengths of frontal and temporal regions were observed in structural networks (with edges weighted by the magnetisation-transfer ratio) of schizophrenia patients compared to controls, and frontal hubs of patients showed a significant reduction of betweenness centrality [14]. Structural networks of schizophrenia patients with auditory visual hallucinations showed higher characteristic path length compared with those of healthy controls [11].

Additionally, studies have shown that schizophrenia patients have higher radial diffusivity and mean diffusivity in some white matter tracts (left thalamo-occipital tracts, right uncinate fascicle, right middle longitudinal fascicle) compared to healthy controls [15, 16]. Another study showed that schizophrenia patients have higher mean diffusivity in the right anterior thalamic radiations, the forceps minor, the bilateral inferior fronto-occipital fasciculus, the left superior longitudinal fasciculus and the bilateral uncinate [54]. Radial diffusivity, although non-specific in principle, can be influenced by myelination and/or axonal density [17, 18, 19], with higher values potentially indicating lower myelination or axonal density.

In this study we used tractography-derived structural brain networks to investigate possible differences in the organisation of the brain networks of healthy participants with varying schizotypy scores. Various microstructural metrics, such as the number of reconstructed streamlines or the fractional anisotropy, have been used in past studies to assign connectivity strength to structural network edges [20, 21, 22]. In this study, in order to capture the possible impact of altered myelination or axonal density on the strength of the connections, we used the inverse radial diffusivity to assign edge weights. The aims of the study were to explore whether previously-reported alterations of white matter connectivity in schizophrenia could also be observed in relation to high schizotypy scores, thereby providing a test of the (dis)continuity between high schizotypy and schizophrenia.

## 2. Methods

Two datasets were collected at the Cardiff University Brain Research Imaging Centre (CUBRIC) between 2014 and 2016 (Cardiff data) and at the University of Munich between 2009 and 2011 (Munich data). The scanners and sequences used were different, and the sensitivity of the data to microstructural parameters is, therefore, different. For that reason, the two datasets were analysed separately.

### Data collection

#### Cardiff data

140 healthy participants (92 female, 19-55 years old, mean=24.7 years, SD=5.0 years) were recruited in and around Cardiff, UK. All procedures were given ethical approval by the Cardiff University School of Psychology Ethics Committee. All participants gave written informed consent before taking part and received financial compensation for their participation. All participants had normal or corrected-to-normal vision and had no history of neurological or neuropsychiatric disorders.

All participants completed the English version of the Schizotypal Personality Questionnaire (SPQ) [23], through which their total schizotypy score was calculated. The schizotypy score is the sum score of 9 subscales: ideas of reference, odd beliefs or magical thinking, unusual perceptual experiences, suspiciousness, excessive social anxiety, no close friends, constricted affect, odd or eccentric behaviour, odd speech. The higher the schizotypy score, the more pronounced the schizotypal traits in a participant. For the questionnaire, adequate internal consistency and criterion validity have been demonstrated [25, 26]. In the Cardiff data, the SPQ subscales showed acceptable to good reliability; the Cronbach’s alpha for the total score was 0.815.

MRI was carried out on a GE Signa HDx 3T scanner (GE Healthcare, Milwaukee, WI). T1-weighted structural data were acquired using an axial three-dimensional fast spoiled gradient recalled sequence with the following parameters: TR = 8 ms, TE = 3 ms, TI = 450 ms; flip angle = 20°; voxel size = 1 mm; field of view (FOV) ranging from 256 × 192 × 160 mm^3^ to 256 × 256 × 256 mm^3^ (anterior-posterior/left-right/superior-inferior). The T1 images were downsampled to 1.5-mm isotropic resolution. Diffusion-weighted MRI data were acquired using a peripherally cardiac-gated sequence with b = 1,200 s/mm^2^, TR = 20 s, TE = 90 ms, voxel size 2.4 × 2.4 × 2.4 mm^3^, zero slice gap, FOV = 230 mm. Data were acquired along 30 unique and isotropically distributed gradient orientations. Three images with no diffusion weighting were also acquired. The diffusion images were co-registered to the T1-weighted images and corrected for head movement and eddy current distortions. Free-water correction was also performed [49, 50].

#### Munich data

115 healthy participants (55 female, aged 18–50 years, mean age = 27.6 years, SD = 8.0 years) were recruited through advertisements in and around Munich, Germany. All experimental procedures were approved by the research ethics committee of the Faculty of Medicine at the University of Munich, in accordance with the current division of the Declaration of Helsinki. Participants were included after thorough clinical screening and only if they did not meet any of the exclusion criteria: any DSM-IV Axis I disorders, first-grade relatives with psychotic disorders, former or current neurological disorders, current physical conditions, current medication except for contraceptives, uncorrected visual impairments. Further inclusion criteria were age between 18 and 55 and German as a first language. All subjects volunteered to take part in the study, gave written informed consent and received financial compensation for their participation.

All participants completed the German version [24] of the SPQ [23] to yield their total schizotypy score. In the Munich data, the SPQ subscales showed acceptable to good reliability; the Cronbach’s alpha for the total score was 0.882.

High-resolution T1-weighted structural images were acquired on a 3T MAGNETOM Verio scanner (Siemens, Erlangen, Germany) using a 12-channel head matrix Rx-coil. A 3-dimensional MPRAGE sequence was used, with repetition time TR = 2400ms, echo time TE = 3.06 ms, flip angle = 9 degrees with 160 slices, slice thickness = 1.0 mm, voxel size = 1.0 × 1.0 × 1.0 mm, field of view FOV = 256 mm. Diffusion-weighted data were acquired using a diffusion-weighted interleaved spin-echo, echo planar imaging sequence with the following parameters: 25 axial slices; slice thickness = 5.2 mm; TE = 107ms; TR = 5s; gap between slices = 5.2 mm; FOV = 230 mm; matrix = 128 × 128; flip angle = 90; bandwidth = 1395 Hz/Px; voxel size = 1.8 × 1.8 × 5.2 mm. Diffusion gradients were applied in 20 directions with b = 1000 s/mm^2^, with each image acquired 3 times. Three images with no diffusion weighting were also acquired.

#### Tractography

Probabilistic, anatomically constrained streamline tractography was performed using MRtrix [27], employing the iFOD2 algorithm [28, 29, 30]. The maximum angle between successive steps was 50 degrees, the minimum and maximum streamline lengths were 30mm and 250mm respectively, and the FOD amplitude cut-off was 0.06. Two million streamlines were generated for each participant, with the seed points on the interface between grey matter and white matter. Visual inspection of the tractograms was performed to verify that the white matter was adequately covered and that streamlines did not extend outside of the white matter.

#### Parcellation and network construction

To validate our analysis against the use of different cortical/subcortical parcellations [58], two atlases were employed to identify the areas of the cerebrum that formed the nodes of the structural brain areas: (i) the AAL atlas [31], which comprises 90 areas, and (ii) the Desikan-Killiany atlas [32], which comprises 84 areas. FreeSurfer (http://surfer.nmr.mgh.harvard.edu) was used to parcellate the T1-weighted anatomical scans [33, 34] and the Desikan-Killiany atlas was used to identify the 84 areas of the cerebrum that formed the nodes of the structural brain networks [27]. The AAL parcellation was calculated using ExploreDTI 4.8.6 [35]. Each dataset was analysed with both atlas parcellations. However, ExploreDTI failed to give a reliable parcellation for the Munich data, possibly due to the lower resolution of the MRI images in that dataset. For that reason, the subsequent analysis for the Munich data was done only using the Desikan-Killiany atlas.

In order to exclude tracts that are not reliable due to having been recovered with a small number of streamlines, we applied a threshold NS_thr_ and excluded edges with fewer than NS_thr_ streamlines. Because that threshold is arbitrary [36], we repeated the analysis for thresholds that correspond to mean network sparsity over participants in the range of 0.85 to 0.25.

Edges that survived thresholding were weighted with the inverse mean radial diffusivity calculated along the voxels spanning the corresponding white-matter tracts. Networks were constructed for the auditory, sensorimotor, visual and default-mode systems, i.e., for subnetworks that have consistently been shown to be altered in schizophrenia patients compared to controls. Three graph theoretical metrics, namely the mean node degree, mean clustering coefficient and mean nodal strength, were calculated for all networks using the Brain Connectivity Toolbox [37]. These metrics provide insight about different aspects of the topological organisation of the brain networks. Specifically, the node degree and the nodal strength measure the connectivity of a node within the network, while the clustering coefficient measures the connectivity within the neighbours of each node. We also note that the node degree is independent of the edge weighting, while the clustering coefficient and the nodal strength do depend on it. In order to identify any graph theoretical metrics that were correlated with each other, correlation coefficients were calculated between graph theoretical metrics for each subnetwork. The mean nodal strength exhibited correlation coefficients with absolute values higher than 0.85 with the mean node degree, and for that reason it was removed from the analysis.

#### Statistical analysis

Correlation coefficients and the corresponding *p*-values were calculated between the demographic characteristics of the participants, i.e., schizotypy score, age and sex, to verify that those were not correlated with each other.

Partial correlation coefficients and the corresponding *p*-values were calculated between the schizotypy score and the graph theoretical metrics of the participants, correcting for age and sex. The mean Cook’s distance [38] was used to identify outliers, which were subsequently investigated further. Data points with Cook’s distance greater than 4 times the mean Cook’s distance were excluded from further analysis, because they either had isolated (unconnected) nodes in their structural connectivity matrices, or were outliers in the degree distribution. The *p*-values that resulted from the correlation analysis were corrected for multiple comparisons using the false-discovery-rate algorithm [39], applied over all graph theoretical measures of all subnetworks.

In order to understand whether any correlations between the graph theoretical metrics and the schizotypy score are due to all the nodes of each subnetwork, or whether they are driven by a few specific nodes of it, we calculated the correlation coefficients between the schizotypy score and the node-specific graph theoretical metrics, for those graph theoretical metrics that exhibited statistically significant correlations. We considered that a node is driving the correlations if the related *p*-value was lower than 0.05.

The analysis pipeline is shown in Fig. 1.

**Fig. 1:**
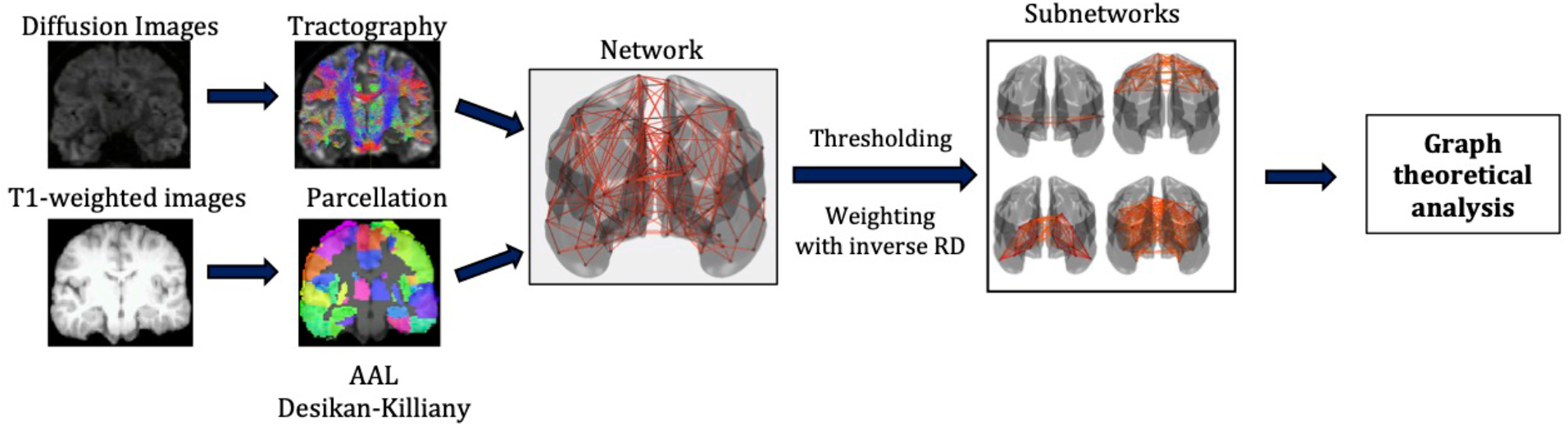
Analysis pipeline.

## 3. Results

### Demographic information

The schizotypy scores (i.e., SPQ total scores) of the participants in the Cardiff data ranged between 0 and 43 (mean: 12.4, SD: 10.1), while for the Munich data they ranged between 0 and 35 (mean: 8.2, SD: 7.0). No statistically significant correlations were found between schizotypy score and age or sex, or between age and sex of the participants, for the Cardiff or the Munich data. The correlation coefficients and *p*-values between these variables are given in Table 1 for the two datasets.

**Table 1:**
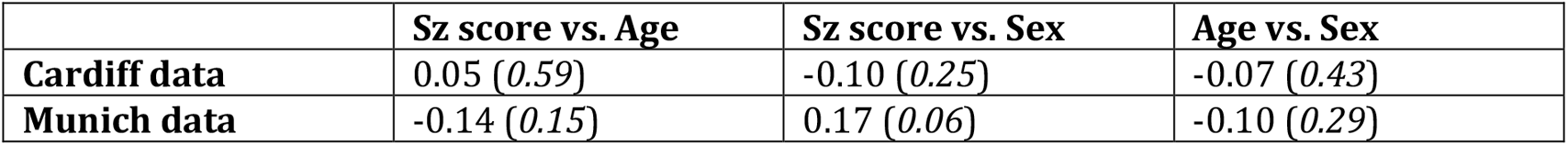
Correlation coefficients and *p*-values (the latter in parentheses) between the demographic data in the two datasets.

### Network analysis

The NS_thr_ required to achieve a given mean sparsity for the whole-brain network in the three cases investigated is shown in Fig. 2. In order to achieve sparsities as low as 0.25, we use NS_thr_ of [0, 160] for the Cardiff/AAL pair, [0, 60] for the Cardiff/Desikan-Killiany pair, and [0, 120] for the Munich/Desikan-Killiany pair. In the following subsections, in addition to showing the correlation coefficients and *p*-values for all these thresholds, we also show scatter plots for the networks that correspond to a sparsity of 0.3, i.e., NS_thr_ = 86 for the Cardiff/AAL case, NS_thr_ = 36 for the Cardiff/Desikan-Killiany case, and NS_thr_ = 55 for the Munich/Desikan-Killiany case.

**Fig. 2:**
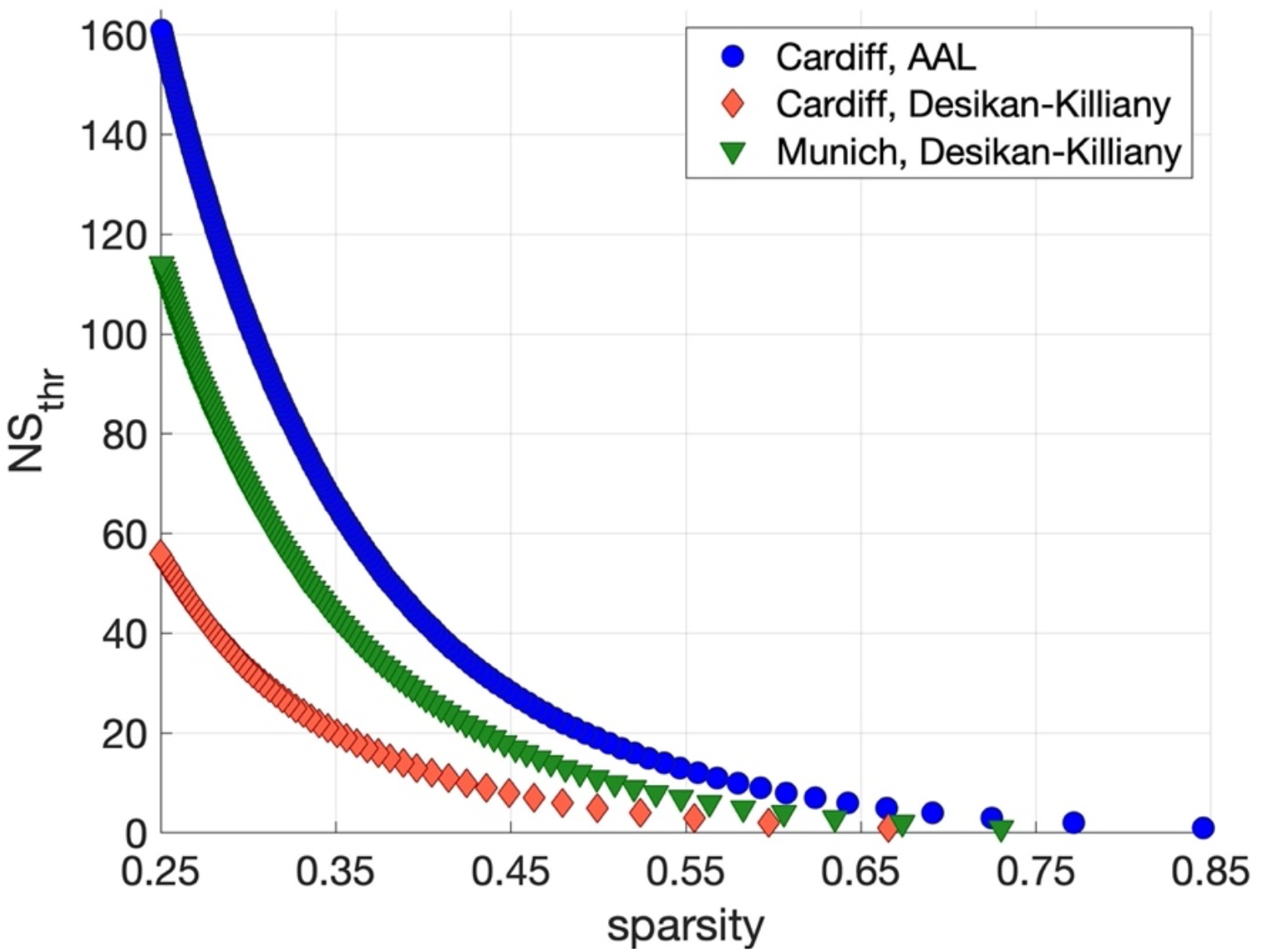
NS_thr_ required to achieve a given sparsity in the whole-brain structural networks, for the three dataset/atlas combinations.

#### Cardiff data, AAL atlas

Significant positive correlations that survived multiple comparison correction and persisted across NS_thr_ were identified between schizotypy score and the mean node degree and mean clustering coefficient, for the sensorimotor and the default-mode networks. The correlation coefficients and *p*-values across NS_thr_ are shown in Fig. 3. Scatter plots for these correlations are shown in Fig. 4.

**Fig. 3:**
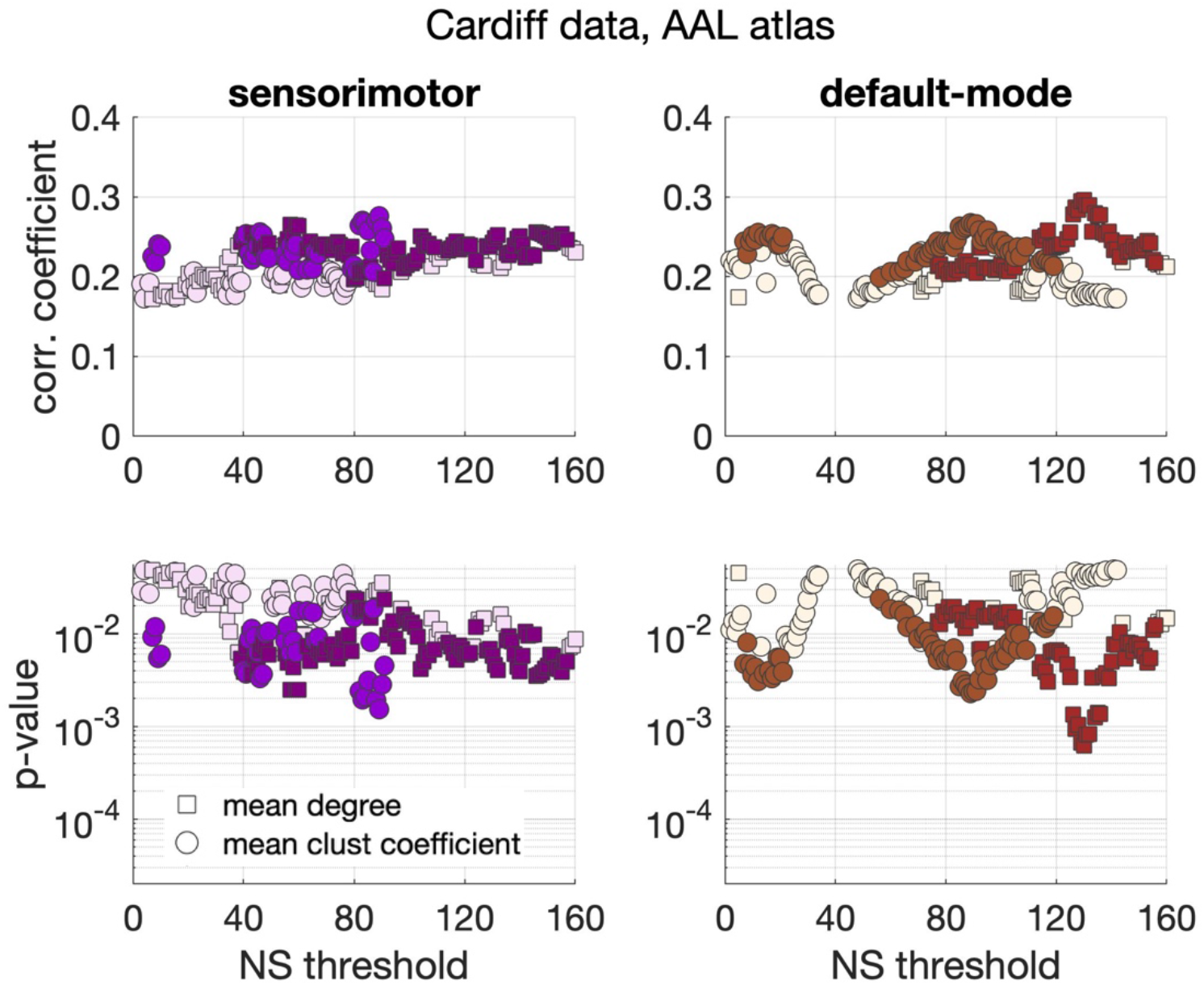
Correlation coefficients and *p*-values between graph theoretical metrics and schizotypy score for the Cardiff data and the AAL atlas. Only the two subnetworks that showed persistent correlations across thresholds are shown. Colored markers indicate correlation coefficients and *p*-values that survived multiple-comparison correction, while blank (fainter colored) markers indicate correlations that were nominally significant (p<0.05) but did not survive multiple comparison correction.

**Fig. 4:**
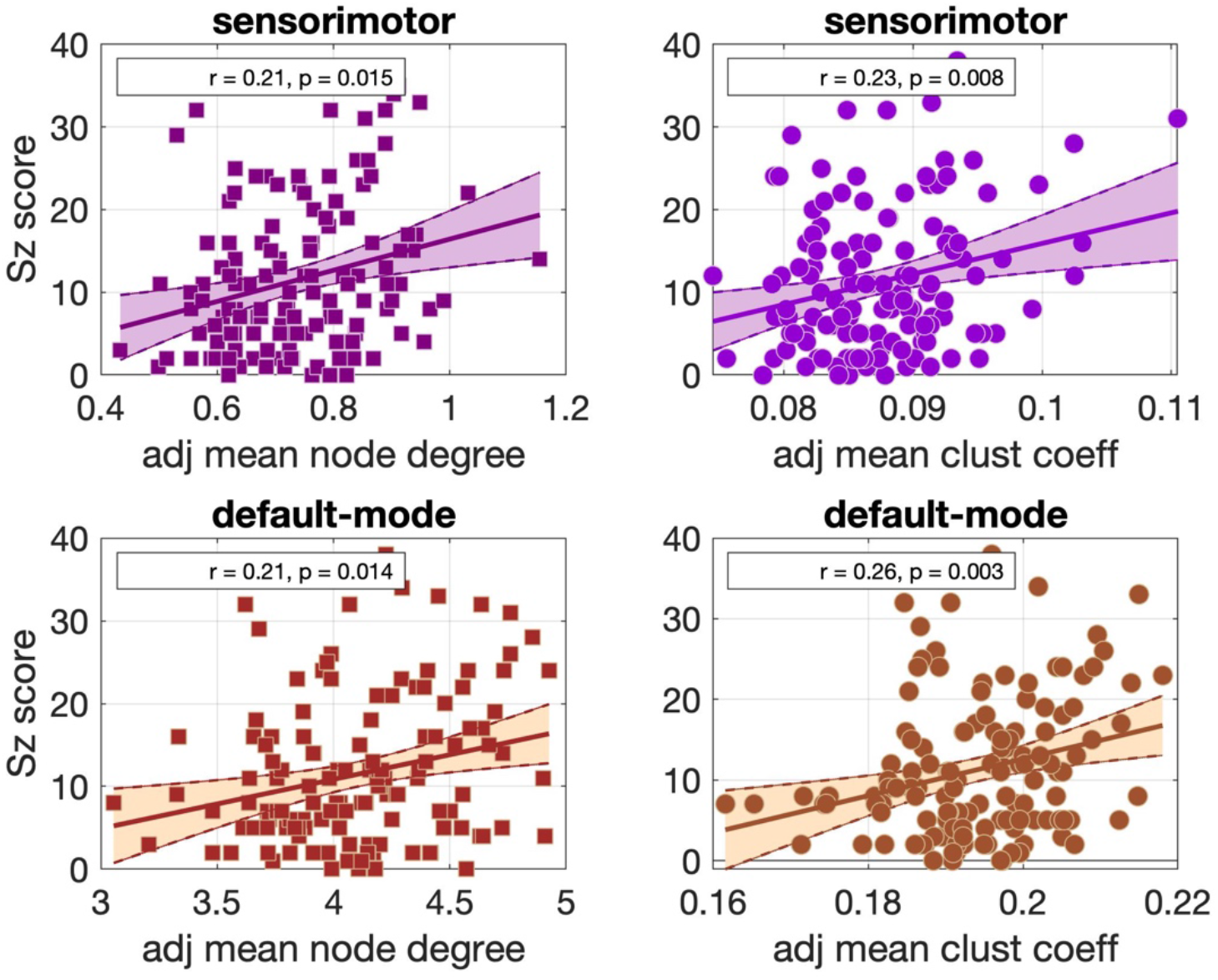
Scatter plots for the correlations observed in the Cardiff data for the AAL atlas parcellation, for NS_thr_ = 86. The best fit line and its 95% confidence interval are also shown.

#### Cardiff data, Desikan-Killiany atlas

Significant positive correlations that survived multiple comparison correction and persisted across NS_thr_ were identified between schizotypy score and the mean node degree of the sensorimotor network, and between the schizotypy score and the mean clustering coefficient of the default-mode network. The related correlation coefficients and *p*-values are shown in Fig. 5. The corresponding scatter plots are shown in Fig. 6.

**Fig. 5:**
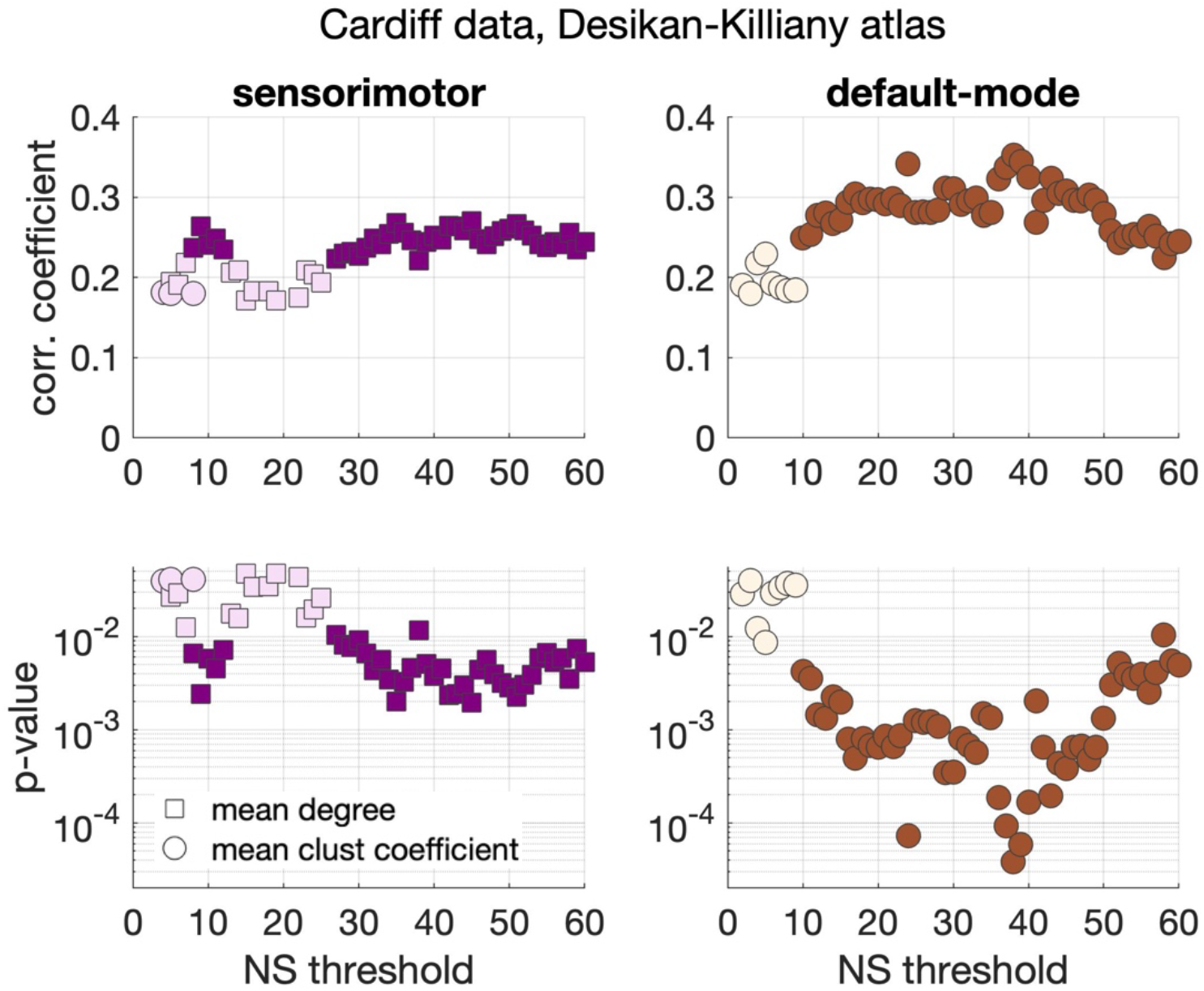
Correlation coefficients and *p*-values between graph theoretical metrics and schizotypy score for the Cardiff data and the Desikan-Killiany atlas. Only the two subnetworks that show persistent correlations across thresholds are shown. Colored markers indicate correlation coefficients and *p*-values that survived multiple-comparison correction, while blank (fainter colored) markers indicate correlations that were nominally significant (p<0.05) but did not survive multiple comparison correction.

**Fig. 6:**
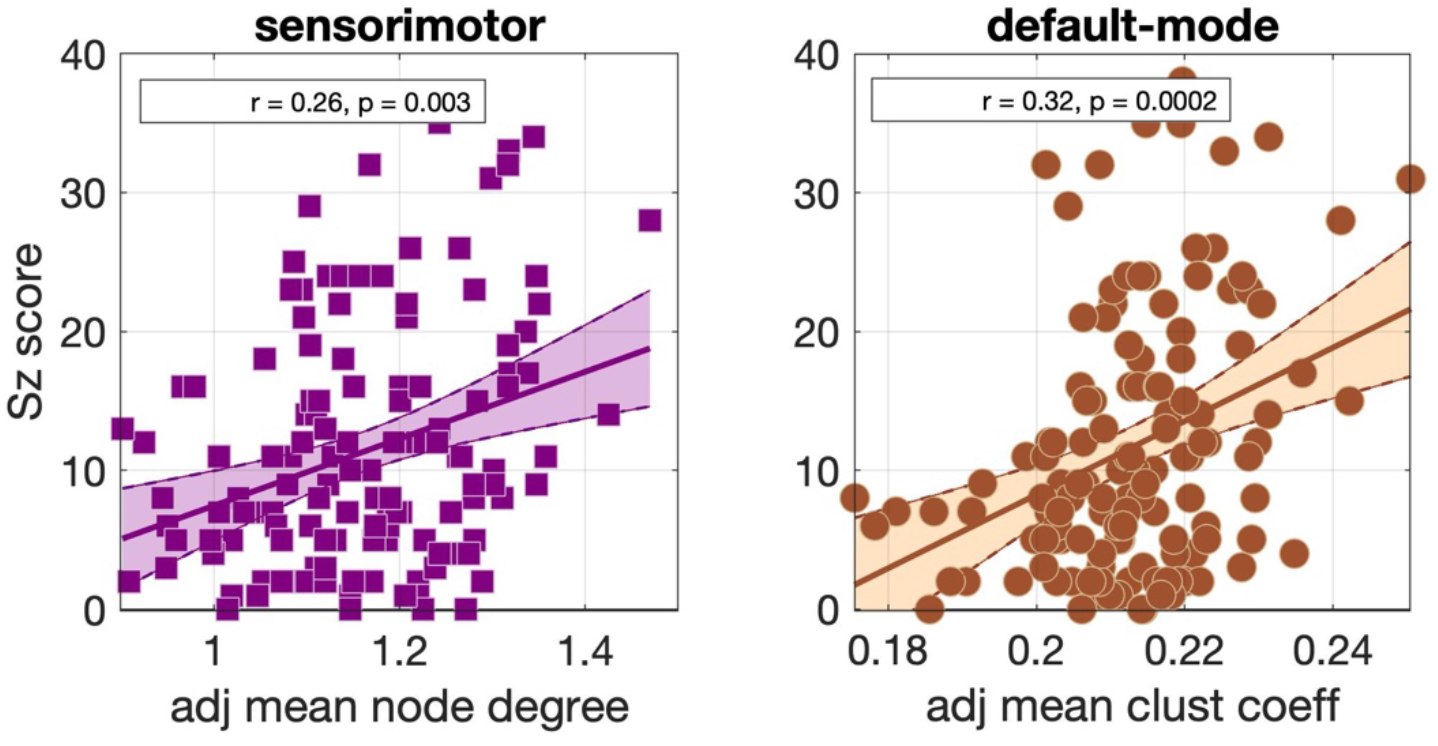
Scatter plots for the correlations observed in the Cardiff data for the Desikan-Killiany parcellation, for NS_thr_ = 36. The best fit line and its 95% confidence interval are also shown.

#### Munich data, Desikan-Killiany atlas

Significant positive correlations were identified between the schizotypy score and the mean node degree of the sensorimotor network, and the mean node degree of the visual network. These correlations were nominally significant, i.e., did not survive multiple comparison correction, but persisted across NS_thr_. The correlation coefficients and *p*-values are shown in Fig. 7. The corresponding scatter plots are shown in Fig. 8.

**Fig. 7:**
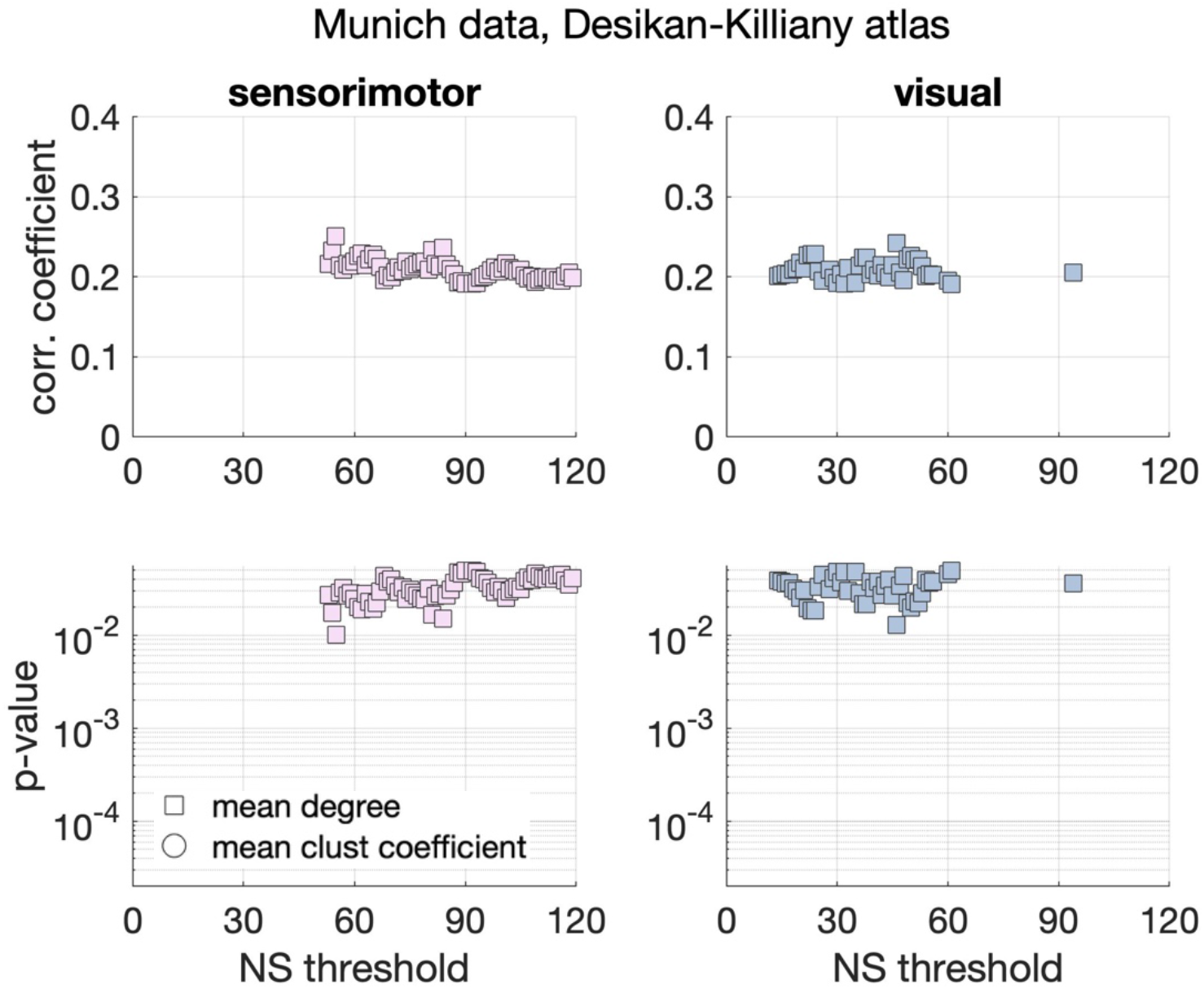
Correlation coefficients and p-values between graph theoretical metrics and schizotypy score for the Munich data and the Desikan-Killiany atlas. Only the sensorimotor and the visual subnetworks showed persistent, nominally significant correlations across thresholds.

**Fig. 8:**
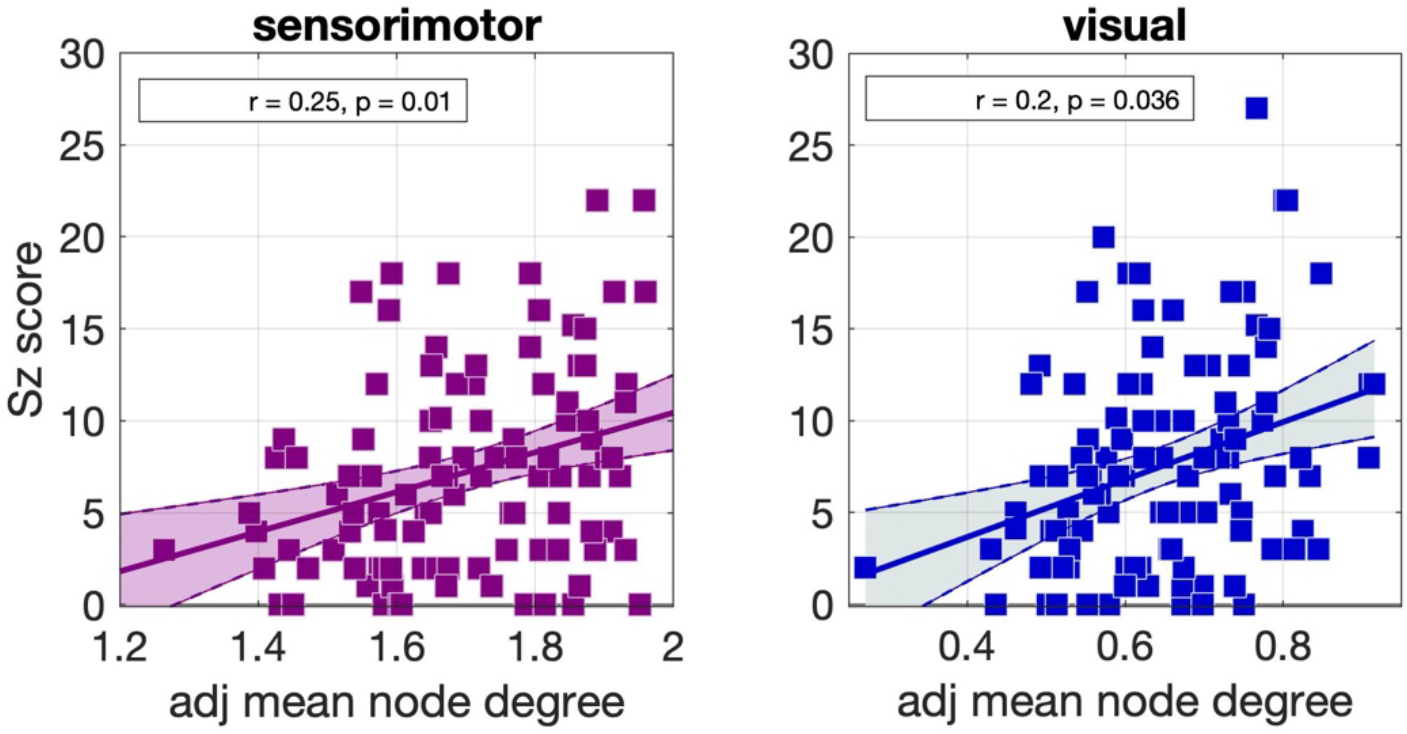
Scatter plots for the correlations observed in the Munich data for the Desikan-Killiany parcellation, for NS_thr_ = 55. The best fit line and its 95% confidence interval are also shown.

### Node-specific analysis

#### Cardiff data, AAL atlas

The node-specific analysis for the sensorimotor network showed that the correlations between the schizotypy score and the mean node degree are driven by the right middle frontal gyrus and the right postcentral gyrus, while the correlations between the schizotypy score and the mean clustering coefficient are driven by the right precentral gyrus and the right postcentral gyrus. For the default-mode network, the correlations between schizotypy score and mean node degree are driven by the left middle frontal gyrus, the right inferior frontal gyrus (opercular part), the left inferior frontal gyrus (orbital and triangular parts), the left and right middle cingulate, the left hippocampus and the left thalamus. Also, for the default-mode network, the correlations between schizotypy score and the mean clustering coefficient were driven by the left and right superior frontal gyrus, the right inferior frontal gyrus (triangular part), the left anterior cingulate, the left and right middle cingulate, the left posterior cingulate, the left parahippocampal area, the middle occipital gyrus, the left angular gyrus and the left and right precuneus. These areas are shown in Fig. 9. Their correlation coefficients and respective *p*-values are given in Table S1 in the Supporting Information section.

**Fig. 9:**
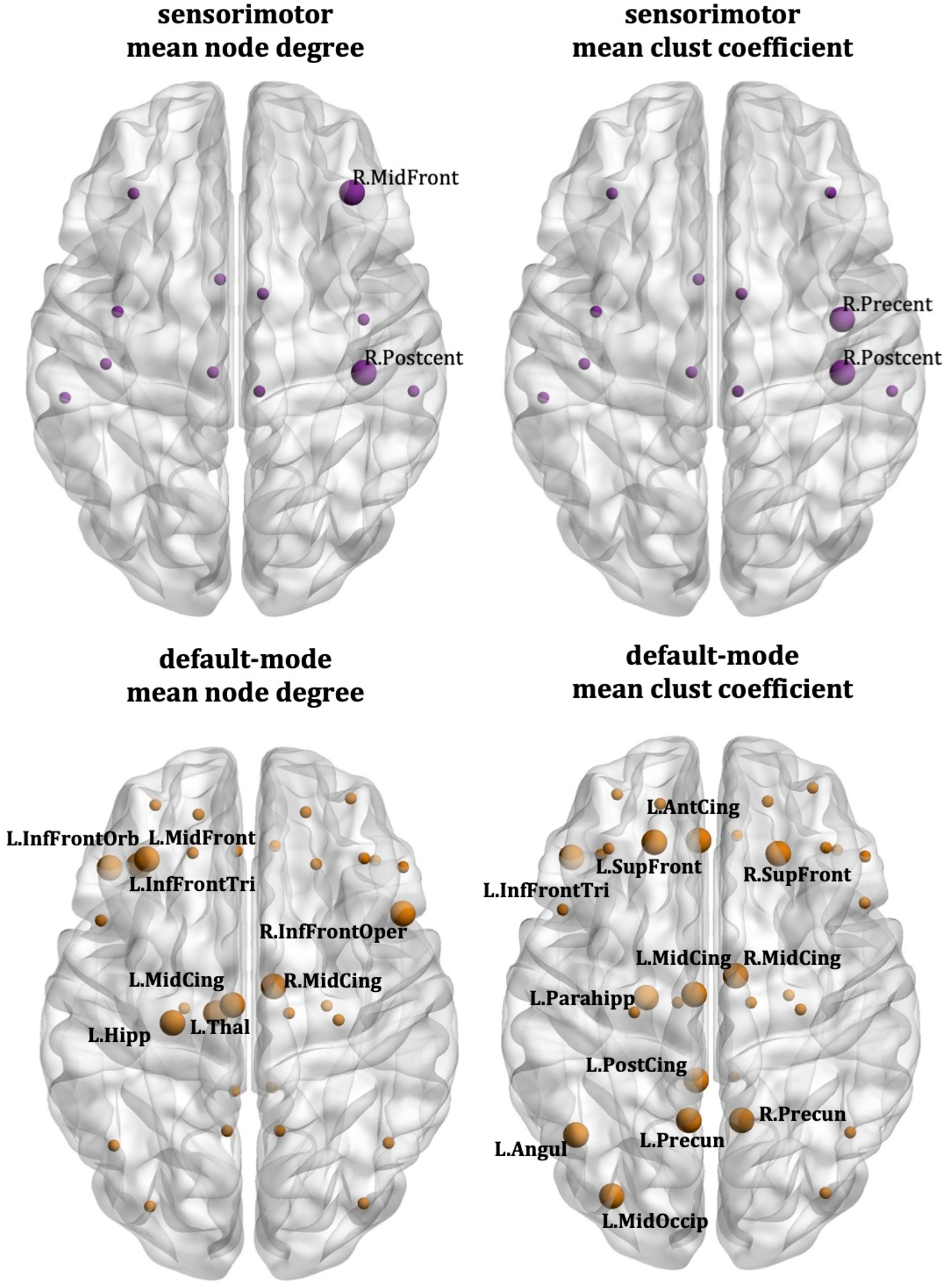
Cardiff data, AAL atlas: Nodes that drive the correlations between the schizotypy score and the sensorimotor mean node degree (top left), sensorimotor mean clustering coefficient (top right), default-mode mean node degree (bottom left) and default-mode mean clustering coefficient (bottom right). The large markers indicate the nodes of each network that drive the correlations; these nodes are also labelled. The small markers indicate the remaining nodes of each network.

#### Cardiff data, Desikan-Killiany atlas

The node-specific analysis showed that the correlations between the schizotypy score and the mean node degree of the sensorimotor network were driven by the left paracentral lobule, the right postcentral gyrus and the right superior frontal gyrus. For the default-mode network, the correlations between schizotypy score and the mean clustering coefficient were driven by the left caudal middle frontal gyrus, the left isthmus cingulate, the left lateral orbitofrontal gyrus, the left and right parahippocampal area, the left and right precuneus, the left and right thalamus, the right hippocampus, the right pars triangularis and the right rostral anterior cingulate. These areas are shown in Fig. 10. Their correlation coefficients and *p*-values shown in Table S1, in the Supporting Information section.

**Fig. 10:**
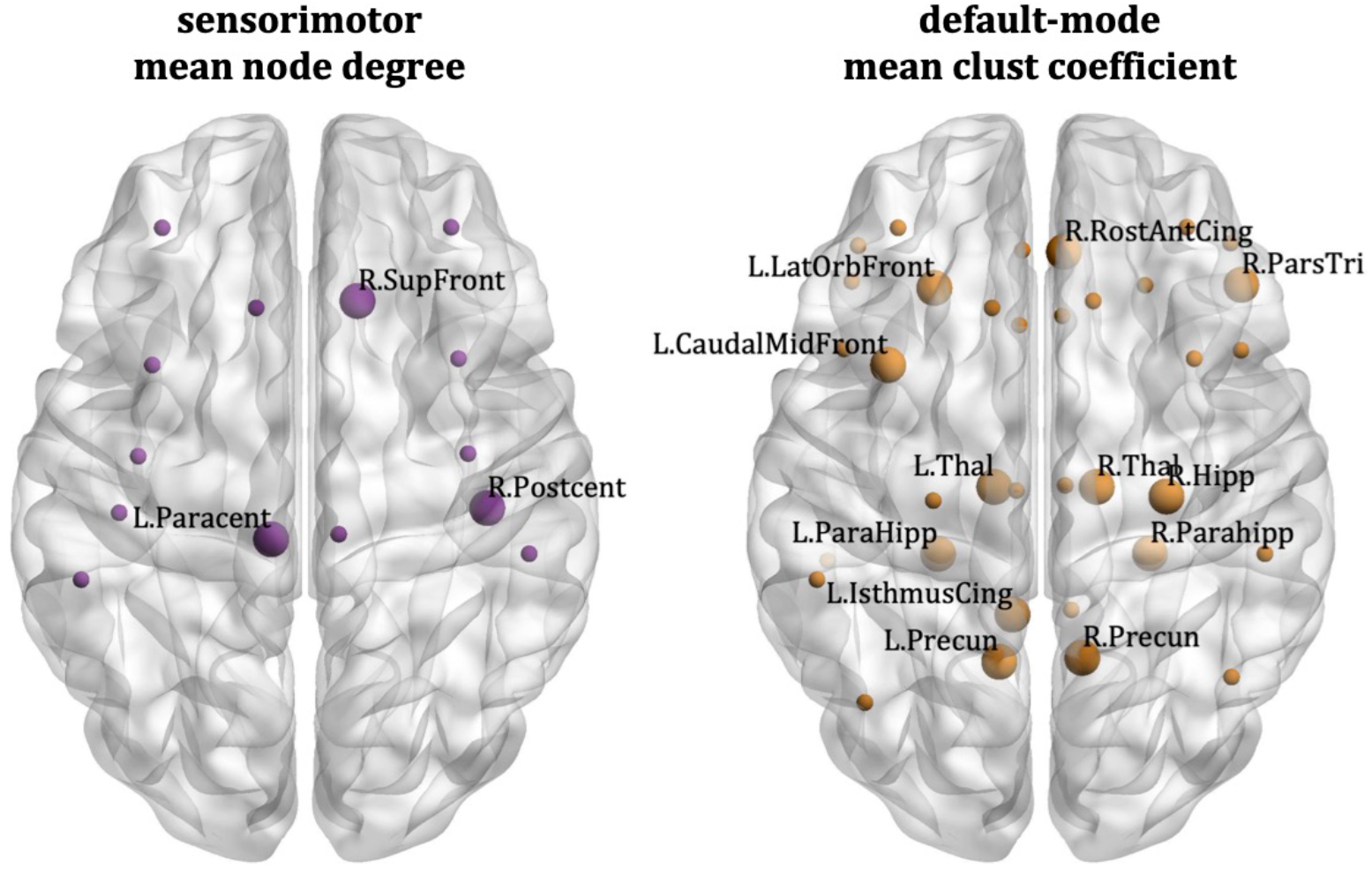
Cardiff data, Desikan-Killiany atlas: Nodes that drive the correlations between the schizotypy score and the sensorimotor mean node degree (left) and default-mode mean clustering coefficient (right). The large markers indicate the nodes of each network that drive the correlations; these nodes are also labelled. The small markers indicate the remaining nodes of the network.

#### Munich data, Desikan-Killiany atlas

The node-specific analysis showed that the correlations between schizotypy score and the mean node degree of the sensorimotor network were driven by the left paracentral lobule, the left and right postcentral gyrus, the left supramarginal gyrus, the right precentral gyrus and the right superior frontal gyrus. The correlations between schizotypy score and the mean node degree of the visual network were driven by the left lingual gyrus, the right cuneus and the right fusiform area. These areas are shown in Fig. 11. Their correlation coefficients and *p*-values shown in Table S1, in the Supporting Information section.

**Fig. 11:**
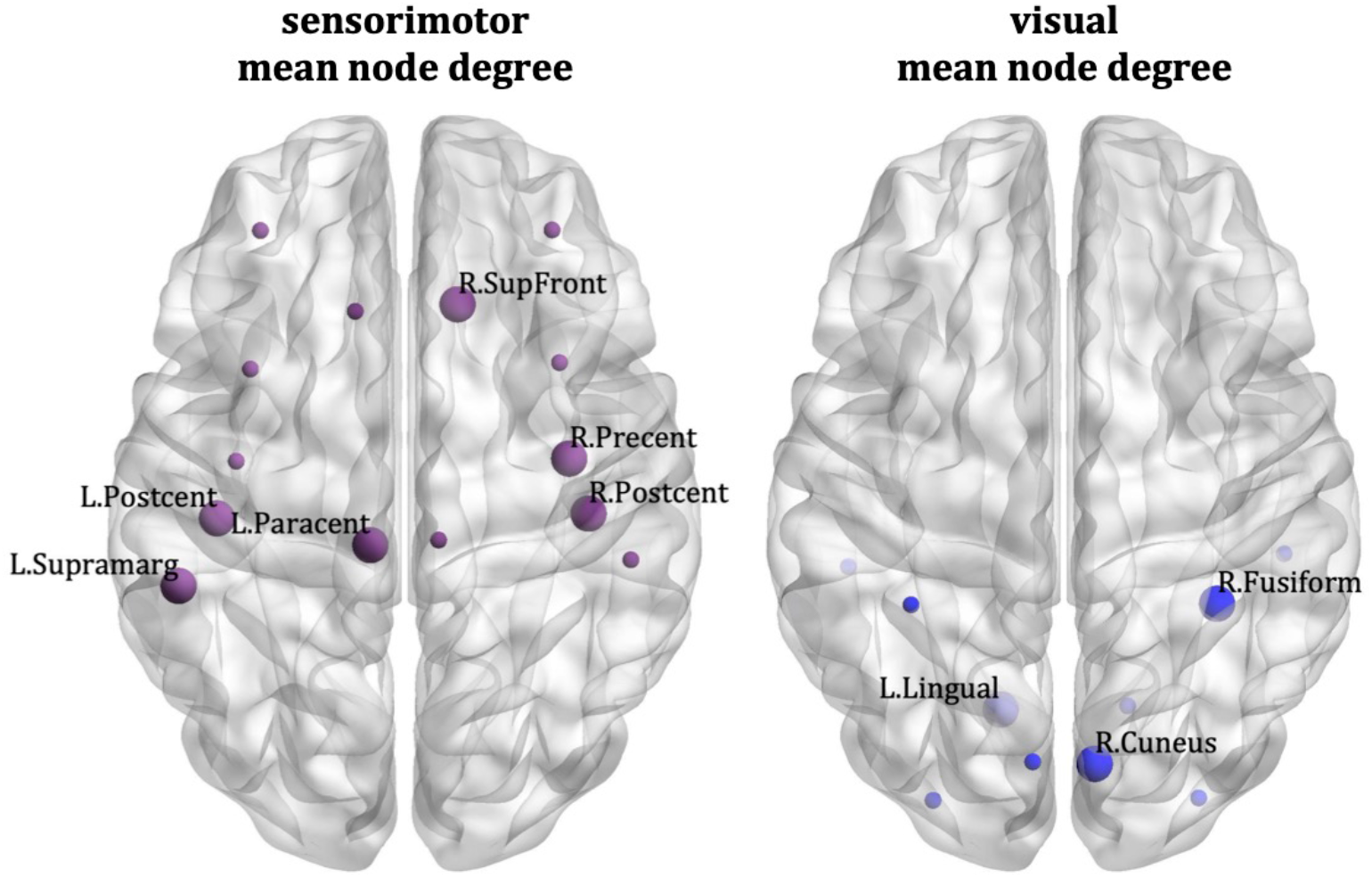
Munich data, Desikan-Killiany atlas: Nodes that drive the correlations between the schizotypy score and the sensorimotor mean node degree (left) and visual mean node degree (right). The large markers indicate the nodes of each network that drive the correlations; these nodes are also labelled. The small markers indicate the remaining nodes of the network.

The visualizations of the nodes were done using BrainNet Viewer [55].

## 4. Discussion

We used graph theoretical metrics to investigate the topology of the structural brain networks of healthy participants with a range of schizotypy scores. To the best of our knowledge, this is the first time these properties are investigated in relation to schizotypy. We used two large datasets, comprising 140 and 115 participants respectively, collected at different locations, in different MRI scanners and with different scanning parameters.

Our results provide evidence that higher levels of schizotypy in healthy participants relate to higher structural connectivity in the sensorimotor and default-mode networks, with some increases also possibly present in the visual network. No evidence of any alterations was found for the auditory network. The edges of the networks were weighted by the inverse radial diffusivity, which partly reflects differences in myelination or axon density [17, 18]. Both myelination and axon density support effective communication in the brain [40, 41]. The edge weighting chosen impacts our results because the clustering coefficient (which exhibited statistically significant correlations with the schizotypy score for the sensorimotor and default-mode network) depends on the values of the edges. On the other hand, the node degree (which also exhibited statistically significant correlations with the schizotypy score) is independent of the edge weights and is only impacted by the number of edges. This means that had we used a different measure to weight the edges (such as, for example the number of streamlines or the fractional anisotropy of the diffusion tensor, which are sometimes used in network studies), the results pertaining to the node degree would have been the same.

The positive correlations observed between the schizotypy score and the graph theoretical metrics of the sensorimotor network were replicated across the two datasets and atlases. Even though they only reached nominal statistical significance in the Munich data, they were persistent across a large range of values of NS_thr_. On the other hand, the positive correlations observed for the default-mode network were not replicated in the Munich data. This could be due to the slightly smaller sample or the lower resolution of the Munich data, or the fact that there were fewer diffusion directions acquired in the diffusion data, as well as a lower b-value – the latter two facts can impact the calculation of the radial diffusivity.

A recent study [42] showed that high schizotypy participants exhibited increased white matter connectivity probability between the right middle frontal gyrus and the right insula, and within the default-mode network. Higher strength values were found between the left superior frontal gyrus and the right rolandic operculum area, and between the right superior frontal gyrus and the right medial superior frontal gyrus [42]. This pattern of higher structural connectivity in high schizotypy agrees with our results. The same study also showed that this enhanced structural connectivity appears alongside lower static functional connectivity in the sensorimotor and default-mode networks in the higher schizotypy participants. While the general pattern of higher white-matter connectivity in relation to higher schizotypy agrees with our finding, we note that differences between the studies include that the study presented in [42] had participants with higher schizotypy scores than the ones in our study, they did not use graph theory in their analysis, and their connectivity measure is more closely related to the number of streamlines rather than to the myelination or to axonal density in the white matter tracts that we used.

Other studies [85, 86, 87] have shown altered microstructural properties in white matter tracts in various schizotypy dimensions. For example, reduced fractional anisotropy (FA) of the cingulum bundle and the uncinate fasciculus was related to symptom severity in schizotypal personality disorder [85, 88]. Also, the FA of the right superior longitudinal fasciculus (SLF) was negatively correlated with disorganised schizotypy [87]. In the same study [87], the axial diffusivity of the right SLF, right anterior thalamic radiation (ATR), right uncinate fasciculus (UF) and forceps minor was positively correlated with negative schizotypy. Finally, the mean diffusivity of the right UF, the right ATR and the forceps minor was positively correlated with negative schizotypy. It should be pointed out that these results pertain to specific white matter tracts, in contrast to our results which pertain to the topological organisation of the entire brain. Those studies also predominantly refer to specific schizotypy dimensions rather than overall schizotypy.

As mentioned in the Introduction, a pattern of lower functional connectivity has been reported in the literature for schizophrenia patients compared to healthy controls, except for some evidence of higher connectivity in the default-mode network. This higher connectivity, however, can be responsible for a failure to completely deactivate the default-mode network during tasks (as happens in healthy participants [72]), which can lead to compromised task performance in schizophrenia. Structural connectivity has also been observed to be lower in schizophrenia, although the findings are less coherent, most likely due to differences in the analysis pipelines used. In this context, there are two possible interpretations for our results.

The first possible interpretation is that our results support the notion that higher structural connectivity (in the presence of higher schizotypy scores) is a protective characteristic against schizophrenia. Specifically, the higher structural connectivity in the sensorimotor network of healthy high-schizotypy participants could be enhancing communication among sensorimotor regions, i.e., regions implicated in action observation and imitation [43], which are functions crucial for successful social interactions. Additionally, the higher structural connectivity in the default-mode network could be leading to more effective communication among regions that are responsible for the emotional engagement during social interactions and the understanding others’ mental states [56], or enhancing self-referential activation [57]. This more effective communication could be protecting these participants from the development of schizophrenia. This interpretation also indicates that structural deficiencies in the sensorimotor and default-mode networks should be further investigated as potential causes of schizophrenia. We note that such potential compensatory effects have been previously reported. For example, increased frontal grey matter and higher frontal activity in schizotypal personality disorder have been suggested as a possible reason for the lower magnitude of cognitive and deficit symptoms in the disorder compared to schizophrenia [80, 81]. At the same time, we also note that schizophrenia is normally diagnosed in the late teenage years to the early 20s in men and in the late 20s to early 30s in women. Some participants in the two samples we used were under 30 years old at the time that the data were acquired. Therefore, some of the higher-schizotypy participants in our samples may, at some time after the data were acquired, develop schizophrenia, which would indicate that the increased structural connectivity in the sensorimotor and default-mode networks could not fully protect them from developing the disease.

A second possible interpretation is that the white matter structural alterations that have been reported in schizophrenia [11, 13, 14, 15, 16] may be the pathophysiological correlate or consequence of worsening schizophrenia symptoms and/or pharmacological treatment, particularly in chronic schizophrenia, rather than the cause of the disease. This is in agreement with the dysconnection hypothesis [44], indicating that functional deficiencies precede structural ones in the disease.

Our findings relating to the nodes of the subnetworks that drive the higher structural connectivity can also be put in the context of existing findings in the literature. We note that there were some similarities and some differences in the nodes driving the correlations for the 3 dataset/atlas pairs. For example, the right postcentral gyrus drove the correlations between schizotypy score and mean node degree for the sensorimotor network in all three cases (Cardiff/AAL, Cardiff/Desikan-Killiany and Munich/Desikan-Killiany), while the left paracentral lobule and right superior frontal gyrus drove those correlations for the later two cases (Cardiff/Desikan-Killiany and Munich/Desikan-Killiany). Additionally, the left parahippocampal gyrus and the left and right precuneus drove the correlations between the schizotypy score and the mean clustering coefficient of the default-mode network for both the Cardiff/AAL and Cardiff/Desikan-Killiany cases. We now discuss the nodes that consistently drove our observed correlations.

The right postcentral gyrus has been shown to have lower functional connectivity with its left counterpart in schizophrenia patients [8]. It has also been shown to have lower grey matter density in schizophrenia patients [45, 46]. Additionally, the global-brain functional connectivity of the postcentral gyrus was lower in adolescent-onset schizophrenia patients compared to controls [47].

The left paracentral lobule was found to have lower regional homogeneity, a quantity that reflects local functional connectivity and integration of information processing in adolescent-onset schizophrenia patients [48]. It was also shown to have lower cortical thickness in psychosis patients compared to healthy participants [70], along with the left pars orbitalis, the right fusiform gyrus and the right inferior temporal gyrus.

The resting-state global-brain functional connectivity (measured via fMRI) of the right superior frontal gyrus was higher in schizophrenia patients and their siblings compared to controls [60]. The right superior frontal gyrus also showed lower grey matter in schizophrenia patients compared to controls [71]. In healthy participants, the grey matter volume of the superior frontal gyrus correlated positively with their SPQ-measured schizotypy scores [78]. Differential functional performance of the right superior frontal gyrus in low vs high schizotypy individuals (albeit measured via the Unusual Experiences subscale of the short form of the Oxford and Liverpool Inventory of Feelings and Experiences, O-LIFE [82]) has also been reported [79], where it was observed that in viewing rejection scenes, low schizotypy participants activated that gyrus, while high schizotypy participants deactivated it.

The higher node degree of these brain areas (right postcentral gyrus, left paracentral lobule and right superior frontal gyrus) within their respective subnetworks (sensorimotor and default-mode) could be responsible for enhanced neural communication in higher schizotypy individuals. This could be protective against some of the functional deficiencies relating to these subnetworks which have been observed in schizophrenia.

Abnormalities in the left parahippocampal gyrus region are a ubiquitous finding in schizophrenia [62]. In an EEG study [59], the left parahippocampal gyrus was found to have higher power spectral density in patients compared to controls. Several structural MRI studies have also reported lower volume in the bilateral parahippocampal gyrus in schizophrenia [63, 64, 65, 66, 67].

The precuneus was shown to have higher connectivity with the posterior cingulate cortex in schizophrenia patients [61]. The resting-state functional connectivity between precuneus and hippocampus was found to have deficits in unmedicated patients with schizophrenia compared to healthy controls [68]. The functional connectivity between the precuneus and the bilateral Heschl’s gyri was found to be abnormal in first-episode schizophrenia [69]. In healthy participants, the schizotypy score was positively associated with gyrification in the precuneus in a study that used the Munich data used in our study [83]. The precuneus is involved in self-processing, mental imagery, episodic memory retrieval, discrimination of self vs others [84]; all these processes might be related to inferring the intention of others, which corroborates the possible role of the precuneus for symptoms within the psychosis spectrum.

The higher clustering coefficient of the left parahippocampal gyrus and the precuneus within the default-mode network implies that their neighbourhoods are better connected in higher schizotypy individuals. This could be compensating for the functional abnormalities observed in these brain areas in schizophrenia. Specifically, if the nodes connected to the precuneus can be used to bypass it, then any functional deficiencies of the precuneus would not affect the brain’s performance in the related functions.

The differences in the nodes that drive the correlations for the different atlases used can be attributed to the fact that the two atlases allocate the cortical and subcortical areas differently, with the AAL atlas comprising 90 cortical and subcortical areas while the Desikan-Killiany atlas comprising 82 cortical and subcortical areas with the remaining two areas corresponding to the two hemispheres of the cerebellum. It has been shown that there are differences in the network sparsity (and therefore the node degree) and in the clustering coefficient as a result of the choice of atlas, e.g., AAL versus Desikan-Killiany, for the same tractography methods [58]. We note, however, that the study presented in [58] used a different edge weight (i.e., connection density) in contrast to the inverse radial diffusivity used in our analysis.

A strength of our study stems from the fact that we used two large datasets, comprising 140 and 115 participants. Despite the differences in scanners, scanning sequences and the language in which the SPQ was administered, the findings of stronger structural connectivity in the sensorimotor network were replicated in the two datasets, and employing two different atlases for brain parcellation. Additionally, the finding of stronger structural connectivity in the default-mode network was replicated when using the two different atlases in the Cardiff dataset. Our tractography generation is state-of-the-art, utilising an algorithm that limits the presence of false positive tracts. Using the inverse radial diffusivity to assign strength to the structural connections relates them to the myelination and the axonal density of the white matter tracts, which are measures that better reflect efficient brain communication than the more frequently used number of streamlines or fractional anisotropy [21]. One limitation of our study is the fact that the schizotypy scores went only up to 43, when the maximum possible value of the SPQ total score is 74. This is a typical range for studies of healthy volunteers but indicates that our results should be interpreted with regards to that population, while generalisations to individuals with very high schizotypy levels need to be drawn with caution. In the future, MRI studies should include dedicated sequences with differential sensitivity to myelin (e.g., magnetization-transfer imaging [73] or myelin-water-based imaging [74]) and to axonal properties, including higher b-value acquisitions, to provide higher sensitivity to the intra-axonal signal fraction and therefore measures such as apparent axon density [75, 76] and axon diameter [77]. This will allow us to further investigate structural connectivity in high schizotypy.

## 5. Conclusions

We investigated the topological properties of structural brain subnetworks in healthy participants of varying schizotypy scores. We found evidence of higher structural connectivity of the sensorimotor and default-mode networks in participants with high schizotypy score. This may indicate a possible protective mechanism against schizophrenia or could provide further support for the hypothesis that functional (rather than structural) dysconnection could be the underlying cause of schizophrenia.

## Acknowledgements

EM was funded by a Wellcome Trust ISSF Postdoctoral Research Fellowship at Cardiff University (204824/Z/16/Z). DKJ was funded by a Wellcome Trust Investigator Award (096646/Z/11/Z) and a Wellcome Trust Strategic Award (104943/Z/14/Z). SF was funded on an Institutional Strategic Support Fund Grant No. 504182 awarded to Cardiff University by the Wellcome Trust. UE acknowledges funding from the Deutsche Forschungsgemeinschaft (ET 31/2-1). This research was funded in whole, or in part, by the Wellcome Trust (204824/Z/16/Z, 096646/Z/11/Z, 104943/Z/14/Z). For the purpose of Open Access, the author has applied a CC BY public copyright licence to any Author Accepted Manuscript version arising from the submission.

## Supporting Information

**Table S1:**
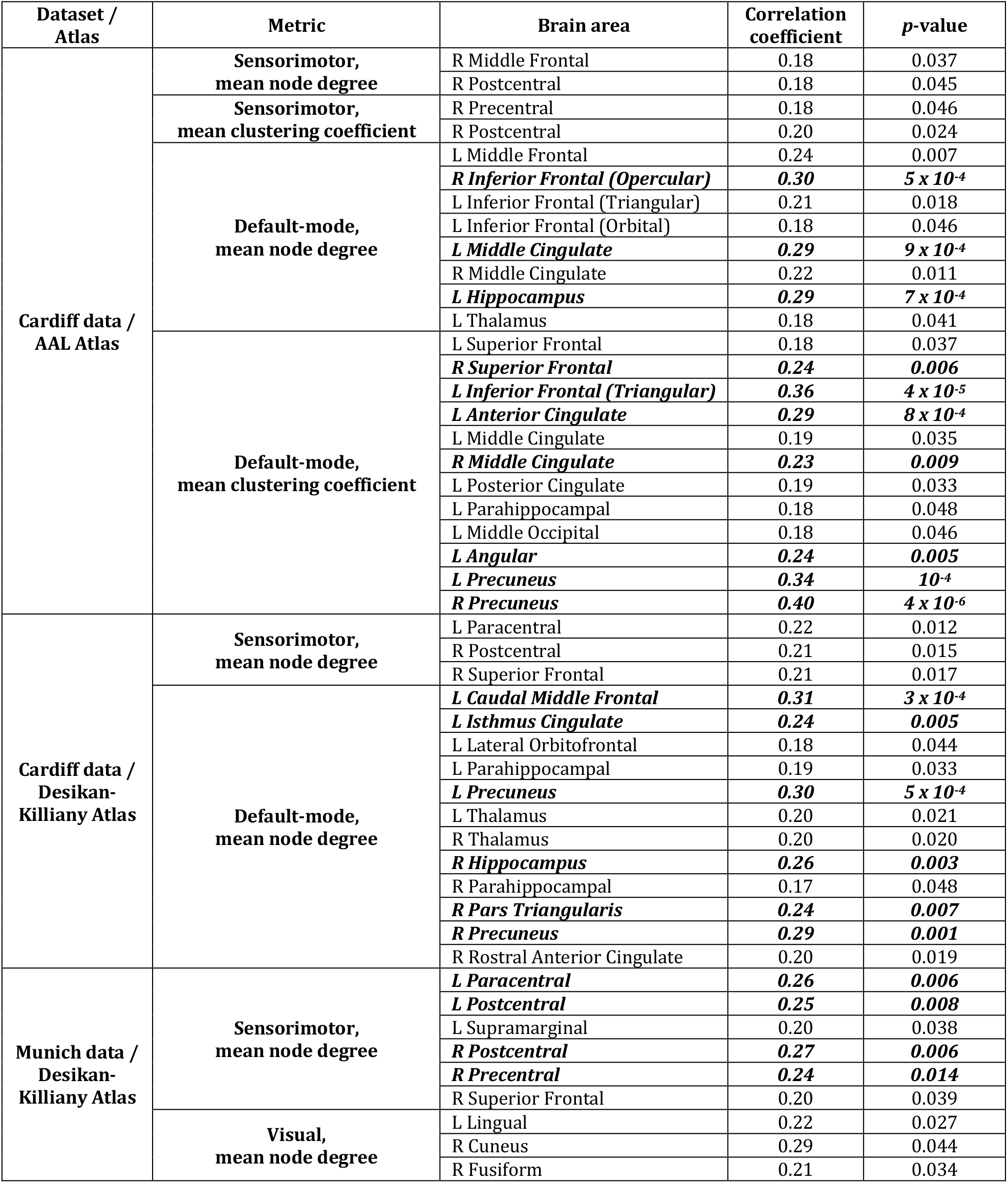
Correlation coefficients and *p*-values between schizotypy score and the graph theoretical measure (node degree or clustering coefficient respectively) of the nodes that drive the correlations. (L = left, R = right). The correlation coefficients and *p*-values that survive multiple comparison correction are denoted in bold italic font.

